# AutoGGN: A Gene Graph Network AutoML Tool for Multi-Omics Research

**DOI:** 10.1101/2021.04.30.442074

**Authors:** Lei Zhang, Ping Li, Wen Shen, Chi Xu, Denghui Liu, Wenjun He, Zhimeng Xu, Chenyi Zhang, Nan Qiao

## Abstract

Omics data identifies biological characteristics from genetic to phenotypic levels during the life span. Molecular interaction networks have a fundamental impact on life activities. Integrating omics data and molecular interaction networks will help researchers delve into comprehensive information underlying the data. Here, we proposed a new multimodal method called AutoGGN to aggregate multi-omics data and molecular interaction networks based on graph convolutional neural networks. We evaluated AutoGGN using two different tasks: cancer type classification and single-cell stage classification. On both tasks, AutoGGN showed better performance compared to other methods, the trend is relevant to the ability of utilizing much more information from biological data. The phenomenon indicated AutoGGN has the potential to incorporate valuable information from molecular interaction networks and multi-omics data effectively. Furthermore, in order to provide a better understanding of the mechanism of prediction results, we assessed the explanation using SHAP module and identified the key genes contributing to the prediction of classification, which will provide insights for the downstream design of biological experiments.

## Introduction

In recent years, high-throughput biomedical technologies such as whole genome sequencing, transcriptome sequencing, Hi-C sequencing and LC-MS-based sequencing, have been widely used in biological research, drug development, and precision medicine^1,2^. Integrating multi-omics data generated from those omics assays, especially comprehensive genomic and transcriptomic data, boosts research and innovation in personalized drug design and precision medication across various research institutions, hospitals and companies^3^. In pharmaceutical companies, multi-omics data are particularly useful in mining potential drug targets and identifying cancer-related genes^4^, which making it indispensable for Research and Development (R&D) process^5^. In the area of precision medicine, integrating gene mutation and expression profiles to identify molecular subtypes of breast cancer will deliver personalized treatment and improve patient care^6^. The integration of multi-omics features can help researchers obtain a comprehensive picture of life development, and establish a deeper understanding of the pathogenesis, development process and molecular mechanisms of diseases.

Deep learning (DL) has demonstrated potential in mining complex and heterogeneous biological data^7^. DL networks such as feedforward fully-connected neural network (FFNN) and randomly-wired residual fully-connected neural network (RRFCN) have already been proven as efficient approaches on omics data analysis^8–10^. These algorithms are powerful for interpreting omics data through fully-connected neural networks, which can form a classification decision from samples with omics features^9,10^.

Meanwhile, molecular networks are also important in understanding the growth of complexity from simplicity in molecular and biomolecular systems^11^. For example, the synthesis of proteases instructed by corresponding genes can catalyze metabolic reactions like lipid degradation^12^. All types of inter and intra-omics interactions form a large and complex biological regulatory network. Therefore, integrating molecular networks and omics data will provide deeper and comprehensive insights into the biological mechanisms associated with the interesting biological problem.

Recently, the rise of graph deep learning provides new insights in analyzing graph data, such as social networks^13^ and consumer-product graphs^14^. Graph-based deep learning (GDL) has been proven to be efficient in inference tasks including node, edge or graph classification, making it one of the hottest topics in machine learning. However, the application of GDL in the biological network and omics data is far less, more researches are needed to dig into the power of GDL^15,16^.

In this paper, we proposed a multimodal method, called AutoGGN, to integrate biological networks and omics data using graph convolutional neural networks. AutoGGN tends to explore the hidden biological patterns behind omics data and biological networks, improving the performance in downstream biological tasks. Specifically, we demonstrated the effectiveness of AutoGGN in but not limited to predicting the cancer subtypes in patients and the development stage of single cells through two example datasets that integrate single/multi-omics data with protein-protein interaction networks, respectively. The contribution of this study is manifested not only in its innovation of integrating omics data with biological networks but also in its effect on enhancing the application of graph-based deep learning in life science fields.

## Results

### Integrating Biological Networks with Omics Data Based on Graph Convolutional Neural Network

Fully-connected neural network showed powerful ability in analyzing omics data and achieved better performance in some tasks^9,10^, but its ability of handle biological networks is limited. AutoGGN, a multimodal method using graph convolutional neural network proposed by us, could integrate molecular interaction networks with omics data (Figure 1) efficiently.

**Figure 1.**
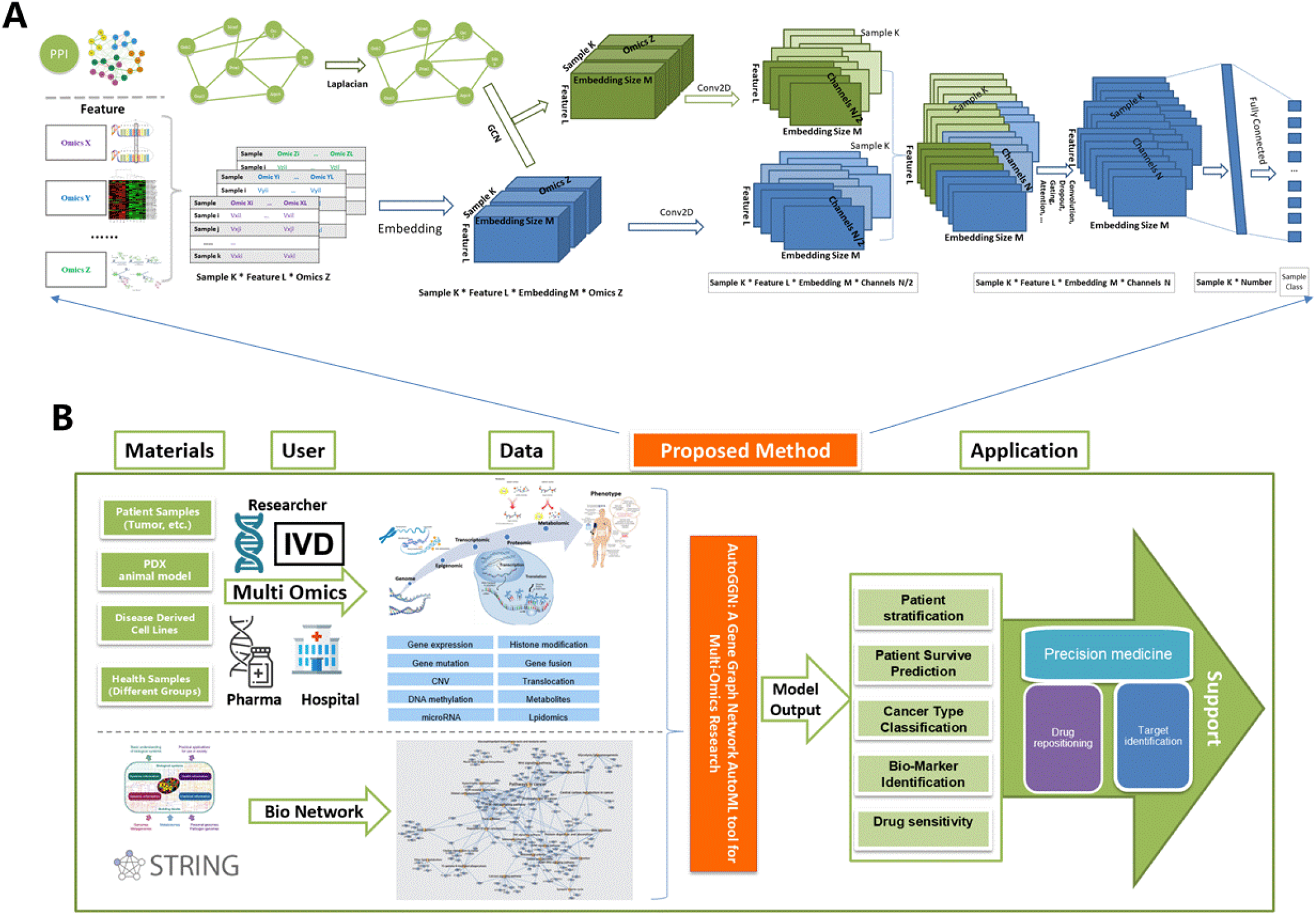
Illustration of algorithm and application for the multimodal method of integrating biological networks and omics Data. (A) Schema of algorithm for the multimodal method of integrating biological networks and omics data based on graph convolutional neural network. The input data includes omics data matrix and molecular interaction network data. Omics data were transformed to high dimension data feature by embedding, then molecular interaction network and omics data would be integrated to fuse the information from biological network into omics data using GCNs. Traditional convolution operator would be performed for both fused matrix and omics feature matrix. Then the results would be concatenated and sent to a fully connected layer. (B) Illustration of application for the algorithm: After researchers get genome/transcriptome/proteomics data from their experimental samples, they can use our innovative model to combine omics data with molecular interaction network. The model can be applied in cancer patient stratification, patient survive prediction, bio-marker identification and drug sensitivity etc.

To get a better representation of omics features, each omics data was mapped to higher dimensions by embedding in AutoGGN. Then different omics channels were concatenated to form a high dimensional multi-channel matrix. For the molecular interactions of omics features obtained from STRING^17^, AutoGGN transformed all the feature interaction pairs to an adjacency matrix. Afterward, graph convolution neural networks^18^ was adopted to convolve the molecular interaction network with the high-dimensional omics data matrix, with the purpose of fusing relation of features and expression of features deeply. After the step, both features of omics data and information of molecular interaction networks were incorporated into the new fusion matrix. Then a traditional convolution operation was performed on the fused matrix and the embedded omics feature matrix separately. The kernel size of the convolution layer was set to 1, intending to merge each feature from different omics separately. Then AutoGGN combined the convolution results from the fused matrix and embedded omics data matrix by channels. It covered the maximum amount of information from omics data and molecular interaction networks. The combined convolution data would be helpful for extracting biological features as well as linking to other operators such as attention layer, gating layer, drop out, and fully connected layer, depending on the type of tasks applied (Figure 1A).

AutoGGN illustrated an innovative approach to integrate molecular interaction networks and multi-omics data through graph convolution neural network. It could also be implemented to solve various biological and medical problems (Figure 1B), including patient stratification, patient survival prediction, biomarker identification and drug sensitivity prediction. In this study, we evaluated AutoGGN on two classification tasks in biology and compared the model performance with other published methods, such as MLP in AutoKeras^19^, RFCN in AutoGenome^9^, AutoOmics^10^. The first case is a cancer type classification task, which involves 9,000 patient samples covering 24 cancer types and two types of omics data (gene mutation and gene expression). The other task is aimed to classify the embryonic developmental stages of mouse single cells. The datasets consist of the gene expression profiles of 1000 single-cell samples covering 10 embryonic stages.

### Validation of AutoGGN on Two Tasks

#### Case 1 – Cancer Type Classification

In the cancer type classification task, 5’780 out of 9’000 patient samples from TCGA^20^ database (https://www.cancer.gov/tcga) had both gene expression and gene mutation data. The patient samples covered 24 cancer types, 5’769 gene features were identified in both gene mutation and gene expression profiles. Based on the protein interaction network obtained from STRING^17^, 260’104 interactions were gotten, including co-expression, co occurrence, gene fusion.

We first applied AutoGGN on the cancer type classification by integrating the protein interaction network and gene expression data of cancer patient samples. A classification model were gotten for 24 cancer type prediction after model training and parameter auto-searching. The performance were evaluated and compare between AutoGGN, XGBoost^21^, AutroKeras^19^ and AutoGenome^9^ by an independent test data, AutoGGN achieved the best accuracy – 0.968, which outperformed XGBoost^21^ (0.911) and AutoKeras^19^ (0.910) by 5 percentage point and AutoGenome^9^ (0.963) by 0.5 percentage point (Figure 2A).

**Figure 2.**
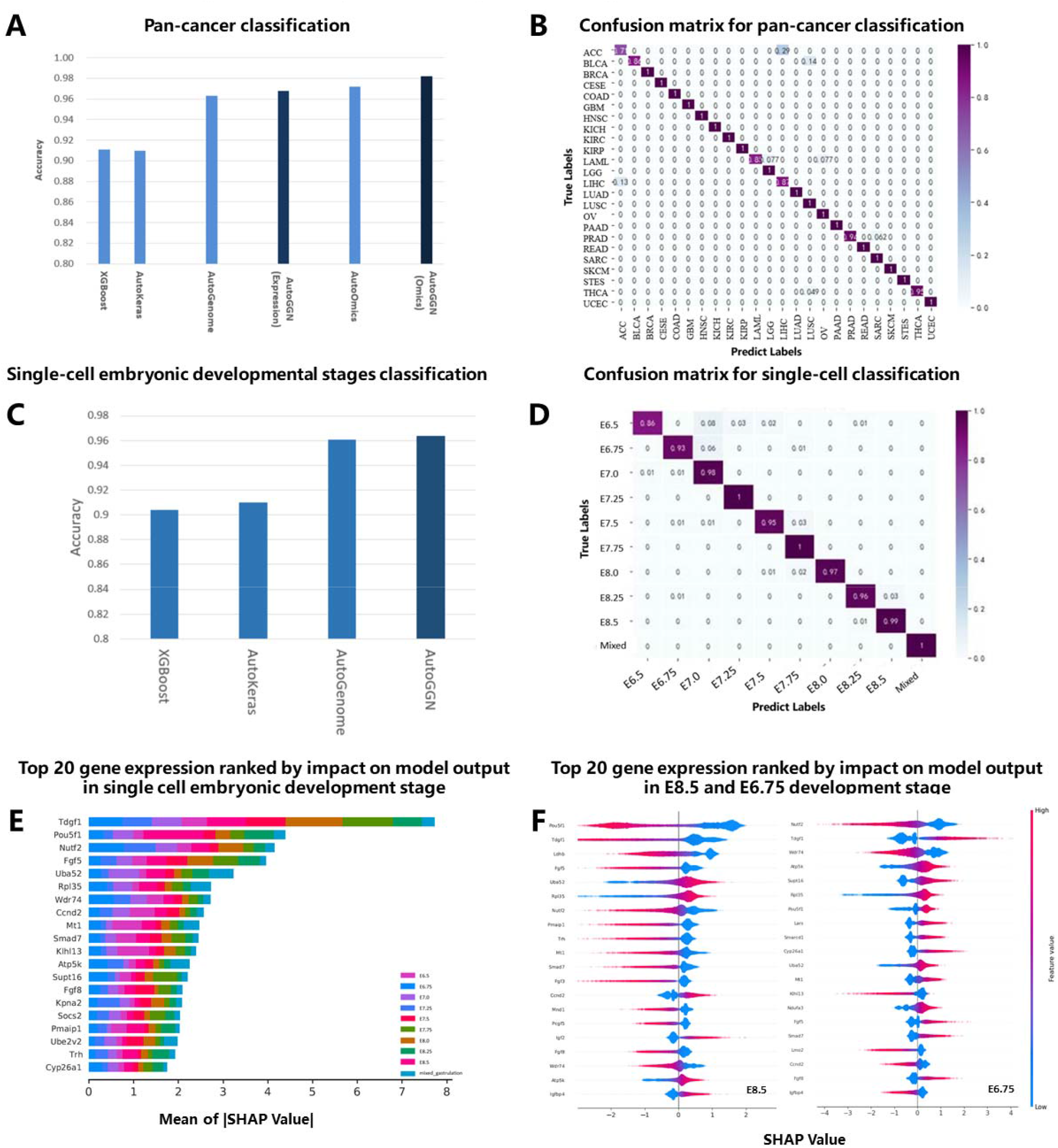
Experimental results for pan-cancer classification and single-cell embryonic developmental stages classification. (A) Accuracy comparison of AutoGGN with XGBoost, AutoKeras, AutoGenome and AutoOmics (Omics) based on cancer type classification. The y-axis indicates the average accuracy score among 24 cancer classification. Single Omic: only gene expression data was used. Omics: both gene expression and gene mutation profiles were used. (B) Confusion matrix for the cancer type classification task. The y-axis indicates the true label, and the x-axis indicates the predicted label. The value in each box was normalized by sample number, indicating the percentage of the samples that had their class correct predicted. The darker purple indicates higher accuracy and the light purple indicates low accuracy. (C) Accuracy comparison of AutoGGN with XGBoost, AutoKeras, AutoGenome based on single cell embryonic developmental stages classification. The y-axis indicates the average accuracy score among 10 different embryonic developmental stages. Single Omic: only gene expression data was used. (D) Confusion matrix for single cell embryonic developmental stage classification. (E) Top-20 genes ranked by feature importance values from gene expression for the 10 single cell embryonic development stage. Colors represents each embryonic development stage. The x-axis indicates the absolute SHAP value. (F) The contribution of the TOP 20 genes expression ranked by SHAP values in E8.5 stage and E6.75 stage. It indicates how gene expression will influence single cell development in different stages. Red represents positive SHAP values that increase the probability of the class, while blue represents negative SHAP values that reduce the probability of the class.

Continually, multiple omics data (both the gene mutation and gene expression data) of patient samples and interaction network were further integrated by AutoGGN. The classification accuracy was 0.982, which was higher than the accuracy 0.968 achieved using single-omics and interaction network. Therefore, AutoGGN outperformed AutoOmics^10^ by one percentage point (Figure 2A), which used multi-omics data as input solely. The detail accuracy for each cancer type was showed in a confusion matrix of AutoGGN (Figure 2B). In general, the classification accuracy was nearly to 1.00, illustrating AutoGGN was able to distinguish the cancer types for most samples with higher resolution.

#### Case 2 – Single-cell stage classification

In single-cell stage classification experiment, a gene expression dataset of mouse single cells covering 10 different cell developmental stages^22^ was used to evaluate performance of AutoGGN. After data preprocessing (Details in Methods), expression profiles of 18’379 genes for 10’000 single cell samples were gotten. Based on the interaction data from STRING^17^, 167,188 protein-protein interaction pairs of high combination score among these genes were collected.

AutoGGN were used to integrate gene expression data and molecular interaction networks of the single-cell samples by graph convolution network. And the performance was assessed by accuracy of classification for single cell stages. Best performance was also achieved by AutoGGN with the accuracy 0.964, which outperformed XGBoost^21^ and AutoKeras^19^ by 5 and 6 percentage point respectively, and was slightly higher than the accuracy of 0.963 achieved by AutoGenome^9^. Confusion matrix (Figure 2D) showed the detailed accuracy for each cell developmental stages predicted by AutoGGN.

Taken together, AutoGGN achieved highest accuracy on both cancer type classification and single-cell stage classification tasks. Model performance is relevant to the degree to which models were able to utilize and analyze biological data effectively. Furthermore, AutoGGN demonstrates the potential of graph convolution neural network on integrating the information from different types of omics data and molecular interaction networks, and achieved a better performance than other models using merely single-omics data or multi-omics data.

### Explain Prediction for AutoGGN using SHAP

Many researchers argued that neural networks has the “black-box” nature^23–25^, which cause poor interpretation of the models despite the high performance they achieve. Explaining and understanding model predictions is equally important besides improving model performance^26^. To alleviated the “black-box” problem brought by deep learning models, we introduced SHapley Additive exPlanations (SHAP)^27^ module into AutoGGN. Given a deep learning model, SHAP would calculate the marginal contribution for each feature to the overall predictions, which was referred as SHAP value. By incorporating SHAP into the model, AutoOmics was able to visualize the SHAP value distributions of each gene to the predicted classes (Figure 2E), and the feature importance of each gene to the predicted classes (Figure 2F). These visualizations could provide meaningful insights towards the deep learning models.

For the single cell embryonic developmental stage classification task, AutoGGN outputted and visualized the top important genes for the classification of developmental stages based on SHAP value (Figure 2E, Figure 2F). We did extensive literature review and found most genes in the top gene list ranked by SHAP value (Figure 2E) were key factors during embryonic development. For example, the protein encoded by Tdgf1, the top 1 ranked gene in the list, is an extracellular, membrane-bound signaling protein that plays an essential role in embryonic development and tumor growth^28^. The top 2 ranked gene – Pou5f1, encodes a transcription factor containing a POU homeodomain that plays a key role in embryonic development and stem cell pluripotency^29^. Another example was the top 4 ranked gene – Fgf5, encodes a protein possessing broad mitogenic and cell survival activities, which is involved in embryonic development^30^. Most of the other genes in the gene list were closely related to embryonic development, proving the predictions of AutoGGN could be clearly interpreted from a biological perspective.

From the SHAP value distributions for each developmental stage, we could also explore how genes contribute to each developmental stage. For example, Pou5f1, Fgf5, Fgf8 and Tdgf1, which were the top ranked genes in developmental stage E8.5 (Figure 2F), were marked in blue for positive SHAP values. It means low expression level of these genes would increase the probability of the E8.5 class. In contrast, high expression level of these genes would increase the probability of an earlier stage E6.75 (Figure 2F). It had been reported that these genes could regulate early embryonic development^28–31^. They were likely to express and regulate activities in early stages of embryonic development like E6.75 and then were controlled under a low expression level in later stages.

## Discussion

In this paper, we proposed AutoGGN, a graph convolutional neural network-based multimodal method, to integrate molecular interaction networks with different types of omics data. This innovative approach is able to utilize and analyze the information from multi-omics profiles and biological network data effectively and comprehensively.

The robustness of AutoGGN was proved in the two classification tasks. In the cancer type classification task, AutoGGN showed its effectiveness in incorporating both single-omics data and multi-omics data with the molecular interaction network. When using gene expression data and interaction network data as input for the model, AutoGGN achieved an accuracy of 0.968, which was much higher than XGBoost^21^ and AutoKeras^19^. When using multi-omics profiles (gene mutation and gene expression) and network data as input, AutoGGN also reached a high accuracy of 0.982 and outperformed AutoOmics^10^ by one percentage point, which was designed to take multi-omics data as input. In single-cell stage classification task, our method demonstrated highest performance among other published models, proving AutoGGN’s capability to make full use of the information from omics features and their interactions during training.

To deal with the black-box challenge of neural networks, we brought SHAP module into AutoGGN. SHAP will function as an explainer for the model output. With an explainer module, AutoGGN is able to visualize the SHAP value distributions as well as the feature importance of each gene to the predicted classes, thus improving researchers’ understandings towards the deep learning models. In single-cell stage classification task, most of the important genes given by the model were closely related to embryonic development. It shows that AutoGGN not only realizes a high performance, but also provide a clear and precise explanation for the model predictions

Besides classification tasks, AutoGGN can also be applied to a wide range of biomedical tasks. For example, our algorithm can be integrated with the Cox proportional hazards model^32^ to predict patient survival. Other applications include bio-marker identification and drug sensitivity prediction. Further work will be continually conducted to extend the AutoGGN model with more applications.

## Methods

### Datasets

In this study, we performed two classification tasks on different datasets to evaluate AutoGGN. Each tasks utilized both single/multi-omics datasets and network datasets. All the datasets involved in our study were downloaded from public websites, and the descriptions of the datasets are detailed below.

#### Pan-cancer dataset (multi-omics)

We downloaded both gene expression profiles and somatic mutation profiles of patients’ tumor samples covering 28 cancer types from The Cancer Genome Atlas (TCGA [https://gdac.broadinstitute.org/]) database^20^.

#### Mouse single-cell transcriptomics dataset (single-omics)

This dataset is the single-cell RNA sequencing data from a timecourse of mouse gastrulation and early organogenesis, which covers 10 different embryonic developmental stages. It was obtained following the instructions provided in https://github.com/MarioniLab/EmbryoTimecourse2018.

#### Protein-protein interaction network

The protein-protein interaction network data were obtained from STRING^17^ database. We used interaction data (protein.links.txt) provided by the database to gather scored links between proteins and accessory data (protein. info) to get the corresponding gene names. Protein-protein interaction networks within Homo sapiens and Mus musculus species were processed separately.

### Data Preprocessing

For both classification tasks, we first applied normalization to the omics data and then selected common gene features between omics dataset and interaction networks. The detailed preprocessing procedures for each case are described as following.

#### Case 1 - Pan-cancer classification

Since several patient samples of 4 cancer types (KIPAN, STAD, GBMLGG and COADREAD) were overlapped with other types, we simply removed these cancer types and used the rest 24 types without overlapping samples to avoid misleading classification. Log2-transformed Transcripts Per Million (TPM) was used to represent gene expression values. We then applied zero-one scaling in a gene-wise manner among the 24 cancer types. Somatic mutation profiles were extracted from TCGA mutation annotation files and represented using 0 for not mutated genes and 1 for mutated genes. After that, the samples were removed if missing either gene expression or mutation profiles. The genes features with missing expression values or mutation information were also removed.

For protein-protein interaction network, we extracted the interactions between genes measured in gene expression and mutation profiles and transformed them into an adjacency matrix utilizing the NetworkX^33^ module in python. The nodes in the adjacency matrix should be consistent with the genes in expression and mutation profiles.

#### Case 2 - Mouse single cell classification

Due to the large sample size of the original dataset (more than 100,000 single cells), we used a subset randomly selected from the original dataset, which consists of 10,000 single cell samples. The expression matrix we utilized contains 22018 genes for 10,000 single cells. These single cells belong to predefined 10 cell types, representing 10 different embryonic developmental stages. Each stage comprises 1,000 cells. The expression matrix was normalized by 0-1 scaling within features before input for model building.

The protein-protein interaction network for mouse single-cell transcriptomics was obtained likewise.

### Model training and evaluation

For the above two tasks, each omics dataset was divided into training, validation and test set with a proportion of 8:1:1. The omics datasets together with the adjacency matrix were input into the model for training. To optimize the model, we applied hyperparameter tuning using random search ^34^ and MBNAS (developed by Huawei) algorithm on the GCN layer number, channel numbers and the embedding size etc. The model achieving highest accuracy for the classification tasks was identified and then evaluated on an independent test set.

### Feature importance estimation

We used the SHapley Additive exPlanations (SHAP) package^27^ to estimate the importance of gene features. The best model identified and the training set were used as input for the SHAP module. Specifically, we used GradientExplainer, an implementation of expected gradients to approximate SHAP values for deep learning models, to conduct the feature importance estimation. After getting the SHAP value of each feature for each sample, we summed the SHAP values of all samples within each class and obtained the importance score of all the features for each class.

## Data Availability

All the data sets utilized in our study are public data. Pan-cancer classification is from TCGA. Single-cell classification data is from accessions: Atlas: E-MTAB-6967 and the processed data is downloaded following the instructions at https://github.com/MarioniLab/EmbryoTimecourse2018. 10X PBMC single-cell RNA-seq was provided by 10X platform and we downloaded the processed expression matrix and cell labels from (https://github.com/ttgump/scDeepCluster/tree/master/scRNA-seq%20data).

## Software Availability

We will open the utilization of AutoGGN package to the public upon the acceptance of manuscript.

## Author Contributions

N.Q. designed and conceived the project. L.Z., P.L. and WS. realized the algorithm of AutoGGN and performed experiments of the project under the guidance of N.Q. C.X., D.L, W.H. and C.Z. discussed and contribute ideas. L.Z., P.L. and W.S. wrote the paper. N.Q. revised the manuscript. All authors read and approved the final manuscript.

## References

1. Fuentes-Pardo, A. P. & Ruzzante, D. E. Whole-genome sequencing approaches for conservation biology: Advantages, limitations and practical recommendations. Mol. Ecol. 26, 5369–5406 (2017).

2. Chen, R. & Snyder, M. Promise of personalized omics to precision medicine. WIREs Syst. Biol. Med. 5, 73–82 (2013).

3. John, A., Qin, B., Kalari, K. R., Wang, L. & Yu, J. Patient-specific multi-omics models and the application in personalized combination therapy. Future Oncol. 16, 1737–1750 (2020).

4. Subramanian I, A. K., Verma S, Kumar S, Jere A. Multi-omics Data Integration, Interpretation, and Its Application. Bioinform Biol Insights 14, (2020).

5. P, T. Big Data in Pharmaceutical R&D: Creating a Sustainable R&D Engine. Pharm. Med 29, 87–92 (2015).

6. Prat, A. et al. Clinical implications of the intrinsic molecular subtypes of breast cancer. The Breast 24, S26–S35 (2015).

7. Xu, S. A., Chunming, Jackson. Machine learning and complex biological data. Genome Biol. 20, (2019).

8. Mahmud, M., Kaiser, M. S., Hussain, A. & Vassanelli, S. Applications of Deep Learning and Reinforcement Learning to Biological Data. IEEE Trans. Neural Netw. Learn. Syst. 29, 2063–2079 (2018).

9. Liu, D. et al. AutoGenome: An AutoML Tool for Genomic Research. bioRxiv (2019) doi:10.1101/842526.

10. Xu, C. et al. AutoOmics: An AutoML Tool for Multi-Omics Research. bioRxiv (2020) doi:10.1101/2020.04.02.021345.

11. Ideker, T., Ozier, O., Schwikowski, B. & Siegel, A. F. Discovering regulatory and signalling circuits in molecular interaction networks. Bioinformatics 18, S233–S240 (2002).

12. Schulze, H., Kolter, T. & Sandhoff, K. Principles of lysosomal membrane degradation: Cellular topology and biochemistry of lysosomal lipid degradation. Biochim. Biophys. Acta BBA - Mol. Cell Res. 1793, 674–683 (2009).

13. Rong, Y. et al. Deep Graph Learning: Foundations, Advances and Applications. in Proceedings of the 26th ACM SIGKDD International Conference on Knowledge Discovery & Data Mining 3555–3556 (Association for Computing Machinery, 2020). doi:10.1145/3394486.3406474.

14. Dr. Adnan A. Ugla, H. J. K., Dhuha J. Kamil. Interpretable Recommender System With Heterogeneous Information: A Geometric Deep Learning Perspective. Int. J. Mech. Prod. Eng. Res. Dev. IJMPERD 10, 2411–2430 (2020).

15. Han, P. et al. GCN-MF: Disease-Gene Association Identification By Graph Convolutional Networks and Matrix Factorization. in Proceedings of the 25th ACM SIGKDD International Conference on Knowledge Discovery & Data Mining 705–713 (Association for Computing Machinery, 2019). doi:10.1145/3292500.3330912.

16. Rossi, E. et al. Temporal Graph Networks for Deep Learning on Dynamic Graphs. (2020).

17. Szklarczyk D, von M. C., Gable AL, Lyon D, Junge A, Wyder S, Huerta-Cepas J, Simonovic M, Doncheva NT, Morris JH, Bork P, Jensen LJ. STRING v11: protein-protein association networks with increased coverage, supporting functional discovery in genome-wide experimental datasets. Nucleic Acids Res 47, (2019).

18. Kipf, T. N. & Welling, M. Semi-Supervised Classification with Graph Convolutional Networks. CoRR abs/1609.02907, (2016).

19. Jin, H., Song, Q. & Hu, X. Auto-Keras: An Efficient Neural Architecture Search System. (2019).

20. Tomczak, K., Czerwińska, P. & Wiznerowicz, M. The Cancer Genome Atlas (TCGA): an immeasurable source of knowledge. Contemp. Oncol. 19, A68–A77 (2015).

21. Chen, T. & Guestrin, C.XGBoost: A Scalable Tree Boosting System. in Proceedings of the 22nd ACM SIGKDD International Conference on Knowledge Discovery and Data Mining 785–794 (ACM, 2016). doi:10.1145/2939672.2939785.

22. Peng, G. et al. Molecular architecture of lineage allocation and tissue organization in early mouse embryo. Nature 572, 528–532 (2019).

23. Kim, B., Khanna, R. & Koyejo, O. O. Examples are not enough, learn to criticize! Criticism for Interpretability. in Advances in Neural Information Processing Systems (eds. Lee, D., Sugiyama, M., Luxburg, U., Guyon, I. & Garnett, R.) vol. 29 2280–2288 (Curran Associates, Inc., 2016).

24. Doshi-Velez, F., Wallace, B. & Adams, R. Graph-Sparse LDA: A Topic Model with Structured Sparsity. (2014).

25. Kim, B., Rudin, C. & Shah, J. The Bayesian Case Model: A Generative Approach for Case-Based Reasoning and Prototype Classification. (2015).

26. Coley, C. W. et al. A graph-convolutional neural network model for the prediction of chemical reactivity. Chem Sci 10, 370–377 (2019).

27. Lundberg, S. M. & Lee, S.-I. A Unified Approach to Interpreting Model Predictions. in Advances in Neural Information Processing Systems 30 (eds. Guyon, I. et al.) 4765–4774 (Curran Associates, Inc., 2017).

28. Strizzi L, et al., Postovit LM, Margaryan NV. Emerging roles of nodal and Cripto-1: from embryogenesis to breast cancer progression. Breast Dis 29, 91–103 (2008).

29. Tantin, D. Oct transcription factors in development and stem cells: insights and mechanisms. Development 140, 2857–2866 (2013).

30. Allerstorfer, S. et al. FGF5 as an oncogenic factor in human glioblastoma multiforme: autocrine and paracrine activities. Oncogene 27, 4180–90 (2008).

31. Abu-Issa, R., Smyth, G., Smoak, I., Yamamura, K. & Meyers, E. N. Fgf8 is required for pharyngeal arch and cardiovascular development in the mouse. Development 129, 4613–4625 (2002).

32. Cai, J. & Zeng, D. Cox Proportional Hazard Model. in Wiley StatsRef: Statistics Reference Online (American Cancer Society, 2014). doi:10.1002/9781118445112.stat06880.

33. Hagberg, A., Swart, P. & S Chult, D. Exploring network structure, dynamics, and function using networkx. https://www.osti.gov/biblio/960616 (2008).

34. Bergstra, J. & Bengio, Y. Random search for hyper-parameter optimization. JMLR 305 (2012).

